# Propagation of genotypic variation across phenotypic levels: a model based on concave relationships unifies disparate observations on phenotypic buffering, dominance, heritability, and inbreeding depression

**DOI:** 10.1101/2025.11.15.688638

**Authors:** Dominique de Vienne, Charlotte Coton

## Abstract

The way genetic variation propagates across successive phenotypic levels up to the fitness components is central to the issue of the genotype-phenotype relationship. The processes involved are highly non-linear, and exhibit a large diversity of behaviors depending on the genes, organization levels, and traits considered. Nevertheless, the shape of the relationship between traits from adjacent levels is predominantly concave, which is probably due to global constraints on matter and energy in the cell. Based on this observation, we used the properties of concave functions to model how phenotypic differences and inheritance vary across increasingly integrated levels of organization. We show that the more integrated the phenotypic levels, the closer the phenotypic values and the larger the positive deviation from additivity (*i*.*e*. dominance or heterosis). These results may explain various observations such as the low heritability and high inbreeding depression of fitness components, and the phenotypic buffering of molecular polymorphisms. Furthermore, the introduction of a cost/crowding factor in the model may explain why overdominance is so rare while heterosis is so common. To our knowledge, this systems biology approach to the genotype-phenotype relationship is the first to be based on a theoretical model of the propagation of genetic variation across phenotypic levels.

## 1 Introduction

In biology, the concept of organization levels and its representation in terms of hierarchical relationships is not merely a practical language tool for conveying knowledge and describing study objects of study. Biological hierarchy is an inherent property of living systems that results from evolutionary processes (Simon, 1962; Szathmáry and Maynard Smith, 1995; Eldredge et al., 2019). It makes these systems intelligible (Bunge, 1977) and raises both epistemological and methodological questions related to emergence, reductionism *vs*. holism and the nature of scientific explanation (Simon, 1962; Korn, 2005).

The familiar “bottom-up” structural organization of increasing complexity — atoms, molecules, macro-molecules, organelles, cells, tissues, organs, organ systems, individuals —, entails several features (Simon, 1962; Campbell, 1974; Umerez, 2016): (i) each level has specific rules and properties, (ii) there are more interactions within levels than between levels and (iii) in addition to the ascending influence of lower levels onto higher levels, there are also descending constraints, sometimes termed “downward causation”, which substantiates the notion of hierarchy.

A central question in quantitative and evolutionary genetics is the relationship between genotype and phenotype, *i*.*e*. what is the relationship between genotypic *differences* and phenotypic *differences* (Orgogozo et al., 2015; de Vienne, 2022). The vast majority of the empirical studies on this topic have focused on the relationship between the genotype and a particular phenotypic level. Very few explicitly address how genetic variation spreads across increasingly integrated organizational levels, up to the fitness components and beyond (populations, communities, ecosystems) (McEntire et al., 2021). Studies that seek to develop formal models that relate variation at a given phenotypic level with variation at higher successive levels are even more scarce. Historically, the pioneer in this field was Wright (1934), who proposed a “Diagram illustrating the chain of processes relating the immediate physiological action of a gene to characters at different levels” (**Figure 1**A), and formalized mathematically the relationship between two of these levels, that of enzymes and that of the metabolic flux. His model revealed that the shape of this relationship is a hyperbola bounded by an asymptote at its upper limit, which results is the dominance of one allele (the “high” allele) over another (the “low” allele). Indeed, the heterozygote value (on the concave hyperbola) is higher than the mean homozygote value (on the line connecting the two parental values). Since then, the genetic and evolutionary implications of the non-linear enzyme-flux relationship have been the subject of a large number of empirical and theoretical studies (reviewed in de Vienne et al., 2023)). Many other models of the transmission of genetic variation between adjacent levels have been proposed, for instance to analyze dominance and epistasis in transcriptional activation cascades (Bost and Veitia, 2014), the relationships between genetic changes in the folding energy of a transcriptional repressor and target gene expression (Li et al., 2019), the homeostasis in the folate-mediated one-carbon metabolism (FOCM) system (Nijhout and Reed, 2014), the molecular epistasis (Domingo et al., 2019), etc. (other examples in Kemble et al., 2019; Alon, 2019).

**Figure 1.**
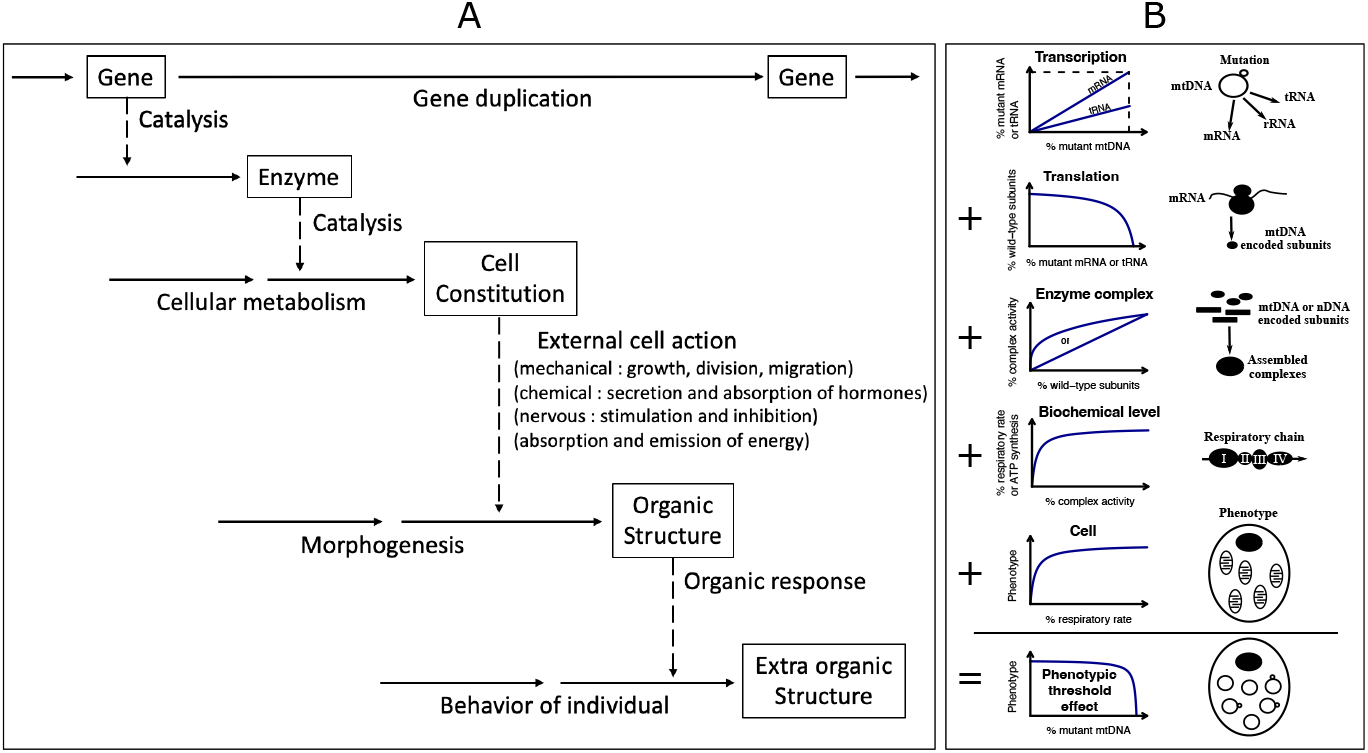
Cascades of phenotypic levels. **A**. Wright’s “diagram illustrating the chain of processes relating the immediate physiological action of a gene to characters at different levels” (redrawn from Figure 2 in Wright, 1934). **B**. Successive relationships between adjacent levels in response to expression of a mtDNA mutation (redrawn from Figure 2 in Rossignol et al., 2003).

However these models only take into account two or three adjacent molecular levels. The propagation of the variation from the genotypic level to the most integrated phenotypic levels still lack general formal approaches. In our opinion, the best example in this direction was given by Rossignol et al. (Rossignol et al., 2003) to explain the “phenotypic threshold effect” observed in the clinical manifestations of mitochondrial genetic defects. The phenotypic effects appear only beyond a certain proportion of mutant mitochondrial DNA (mtDNA), which was explained qualitatively by the non-linear, concave relationship observed at the successive transitions: mRNA/tRNA → enzyme subunits → enzyme complex activity → mitochondrial respiration/ATP synthesis → cell growth → clinical features (**Figure 1**B).

This particular case raises the question of whether it is possible to develop a theoretical framework, or at least identify general principles, to describe the propagation of genotypic variation across increasingly integrated phenotypic levels. This goal may seem completely unrealistic, because the relationship between two adjacent or distant levels can take many non-linear forms, *e*.*g*. gradual responses, sigmoid functions, step functions, increasing then decreasing functions, bimodal responses, chair-shaped relationships, etc. (Pryciak, 2008; Nijhout and Reed, 2014; Keren et al., 2016; Kemble et al., 2019), due to the large diversity of molecular mechanisms involved (Kauffman, 1993; Domingo et al., 2019; Alon, 2019). However, it is worth noting that two types of relationship are predominant:

i. Diminishing returns” relationships. As the activity or abundance of a given factor increases (other factors remaining constant), the output decreases, resulting in a concave response curves reaching saturation. In addition to the case of enzyme-flux relationships, which generally exhibit this behavior, and the case of the mtDNA phenotypic threshold effect mentioned above, diminishing returns have been observed for various transitions between phenotypic levels, such as the transition between transcription or translation machinery factors and gene expression level (Giorgetti et al., 2010), protein synthesis rates (Firczuk et al., 2013), morphological traits (Rosas et al., 2010), and fitness (MacLean et al., 2010; MacLean, 2010; Tokuriki et al., 2012). Actually, the law of diminishing returns could be a general property of the matter-energy exchange networks, as suggested by Petrizzelli et al. (2024) from their formal analogy between electrical circuits and metabolic networks.
ii. Sigmoidal (S-shaped) relationships. Such relationships are commonly observed as a response of the transcription rate to an increase in the concentration of an activator. Cooperativity (Veitia, 2003; Kazemian et al., 2013) and synergy (Carey, 1998) are the main mechanisms underlying this type of response, which can act as a molecular switch, with thresholds and boundaries (Gibson, 1996; Perry et al., 2012; Li et al., 2019). S-shaped relationships can also be observed at other levels, for example as a response to the genetic variations of protein kinase (MAPK) cascade components (Ferrell Jr, 1997; Nijhout et al., 2003), and between remote levels as shown in yeast in the relationship between the expression of *LCB2*, a gene involved in sphingolipid synthesis, and fitness (Rest et al., 2013). Note that for selective reasons, it is likely that most of the alleles in natural populations tend to be positioned on or close to the plateau of the sigmoidal curves, or at least beyond the inflection point (Tokuriki and Tawfik, 2009; Domingo et al., 2019), *i*.*e*. in the concave part of the curve.

Thus, concavity, whether it corresponds to the right-hand side of the sigmoidal curve or to a diminishing returns response, seems to be the most widespread type of relationship between phenotypic levels. Therefore, although it is not possible to develop general models for the propagation of genetic variation across phenotypic levels, it is relevant to try to model the most common type where relationships between successive phenotypic levels are concave.

Using the general properties of concave functions, we show that, as phenotypic levels become more integrated, phenotypic differences between individuals fade while deviation from additivity in crosses increases. These results are valid irrespective of the concave functions involved and fully congruent with various observations from quantitative genetics, such as the lower heritability and higher inbreeding depression of fitness components compared to less integrated traits, and phenotypic buffering at higher levels of biological organization.

## 2 Model of concave genotype-phenotype relationships

### 2.1 Theoretical background

We consider the relationship between the DNA level (the genotype) and increasingly integrated phenotypic levels: transcription rate, enzyme activity, metabolic fluxes, …, up to fitness. We have the following cascade:

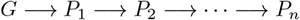

where *G* is the genotype and *P*_*i*_ is the *i*^th^ phenotypic level for all *i* ∈ [1, *n*]. Traits are quantitative and can be monogenic or polygenic.

#### Monogenic traits

Consider a biallelic genetic variation (*A*_1_*/A*_2_), with three genotypes *A*_1_*A*_1_, *A*_1_*A*_2_ and *A*_2_*A*_2_. Following Falconer and Mackay (1996), the genotypic variable *x* is the number of doses of allele *A*_2_. Thus we have

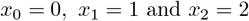

The phenotypic value at level *P*_1_ is a function of *x*:

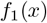

The value at level *P*_2_ is function of the value at level *P*_1_:

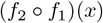

and so on. Thus, the relationship between the genotypic level and the phenotypic level *P*_*i*_ is given by the composition of functions

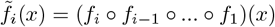

We defined the functions *f*_*i*_ between *P*_*i−*1_ and *P*_*i*_ as monotonically increasing concave functions (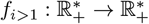 and 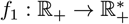).

The phenotypic values of the three genotypes *A*_1_*A*_1_, *A*_1_*A*_2_ and *A*_2_*A*_2_ at level *P*_*i*_ are respectively

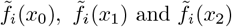

The heterozygote value 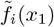 can be expressed in terms of the homozygote values by using the formalism of concave functions:

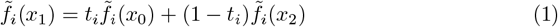

where *t*_*i*_ ∈ [0, 1] if we assume that the heterozygote value lies between the homozygote values. Note that *t*_*i*_, the *deviation from additivity* at level *P*_*i*_, is strictly equivalent to the dominance coefficient *h* used in evolutionary genetics (Crow and Kimura, 1970) and to the degree of dominance *D* first defined by Wright (1934). Because the functions *f*_*i*_ are all increasing concave functions, the heterozygote value is higher than the homozygote mean and does not exceed the highest homozygote value. Thus *t*_*i*_ ∈]0, 0.5[, *i*.*e*. there is partial dominance. At the genotypic level *G* we have *t*_0_ = 0.5 because 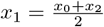.

#### Polygenic traits

Consider now a polygenic trait whose variation depends on *n* biallelic loci, each one controlling a component of the trait. For example, the trait is a metabolic flux controlled by *n* genes, each affecting the concentration or activity of an enzyme in a pathway.

The phenotypic values of component *j* for the three genotypes at level *P*_*i*_ are noted

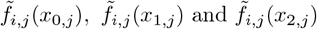

with

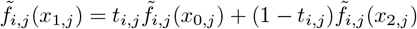

where 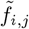 is the composition of functions relating *G* to *P*_*i*_ for component *j* and *t*_*i,j*_ is the deviation from additivity of component *j* at level *i*.

The value *z*_*i*_ of a polygenic trait at level *i* is assumed to be an *n*-dimensional increasing concave function *g* of all the components. Thus we have:

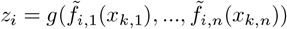

where *x*_*k,j*_ takes the value 0, 1 or 2 depending on the genotype at the locus controlling component *j*.

In the following, we consider two lines, L1 and L2, and their hybrid L1xL2. The genotypes of L1 and L2 are an assortment of “high” and “low” alleles. For instance:

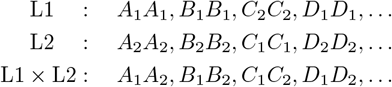

The subscript 1 corresponds to low alleles and 2 to high alleles. By convention, the phenotypic value of L2 is higher than that of L1: *z*_*i*,L2_ > *z*_*i*,L1_. The phenotypic value of the hybrid is

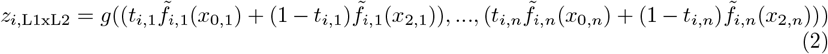

All symbols used are listed in **Appendix A**.

#### Questions addressed

Assuming concavity for every transition *G → P*_1_ and *P*_*i*_ *→ P*_*i*+1_, we addressed two questions:

1. How do the phenotypic differences between individuals change across successive phenotypic levels?
2. How does the inheritance change across successive phenotypic levels?

### 2.2 Phenotypic values converge as phenotypic levels become more integrated

From one phenotypic level to another, the traits considered may be expressed in terms of different units of measure (*e*.*g*. a metabolic flux in mol.L^*−*1^.s^*−*1^, a growth rate in cm.day^*−*1^ and the grain yield in t.ha^*−*1^). Therefore, in order to compare differences between individuals from one phenotypic level to another, we divided the highest homozygote value by the lowest to obtain dimensionless ratios of phenotypic values.

#### Monogenic traits

For monogenic traits, we compared:

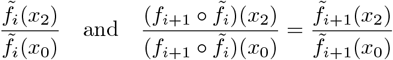

These ratios were noted *q*_*i*_ and *q*_*i*+1_, respectively. In **Appendix B**, we show that

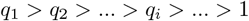

which means that the homozygote phenotypic values converge as phenotypic levels become more integrated, irrespective of the type of concave function.

To visualize this reduction in phenotypic distance, we considered a sequence of four levels. Each transition (*G → P*_1_, *P*_1_ → *P*_2_, *P*_2_ → *P*_3_, and *P*_3_ → *P*_4_) was modeled with a hyperbolic function of the form 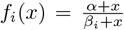, with arbitrary *β*_*i*_-values to obtain different degrees of curvature. The increase in curvature with increasing phenotypic integration is shown in **Figure 2**A, and the related decrease of *q*_*i*_ is shown in **Figure 2**B.

**Figure 2.**
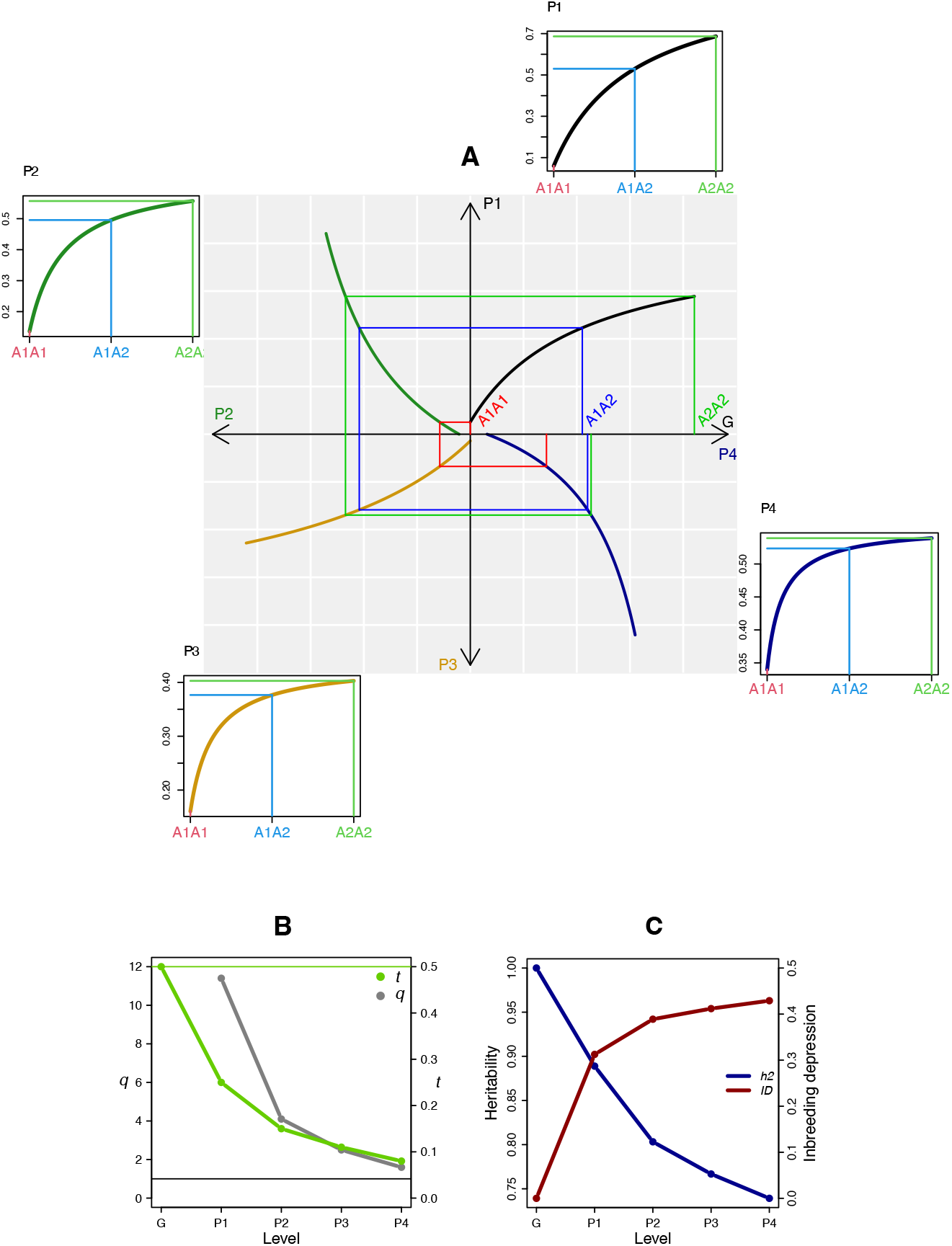
Phenotypic convergence and deviation from additivity across the phenotypic levels for a monogenic trait. **A**. Graph of arbitrary concave relationships between the genotypic level *G* and four successive phenotypic levels, presented as four sub-graphs rotating counter-clockwise on the main graph (light gray background). Top right sub-graph: *G → P*_1_ relationship 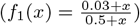. Top left sub-graph: *P*_1_ *→ P*_2_ relationship 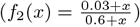. Bottom left sub-graph: *P*_2_ *→ P*_3_ relationship 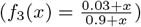. Bottom right sub-graph: *P*_3_ *→ P*_4_ relationship 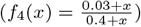. The genotypic values are *x*_0_ = 0 (homozygote *A*_1_*A*_1_, in red), *x*_1_ = 1 (heterozygote *A*_1_*A*_2_, in blue) and *x*_2_ = 2 (homozygote *A*_2_*A*_2_, in green). The evolution of the red, blue and green lines from *G* to *P*_4_ show that parental values become closer to each other and there is increasing deviation from additivity. The four insets next to the sub-graphs show the increasing curvature from the relationships *G → P*_1_ to *G → P*_4_. **B**. The increase in curvature results in a decrease of both *q*_*i*_, the ratio of homozygote phenotypic values, and *t*_*i*_, the deviation from additivity. **C**. As phenotypic levels become more integrated, narrow-sense heritability decreases (blue curve) and inbreeding depression increases (red curve).

#### Polygenic traits

For polygenic traits, the ratio 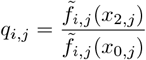 of every component *j* decreases and converges to 1 with increasing phenotypic integration. Because the function *g* is continuously increasing in all dimensions, the more integrated the polygenic trait, the closer its ratio 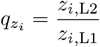 is to 1. Thus

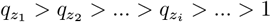

### 2.3 Deviation from additivity increases with phenotypic integration

#### Monogenic traits

The heterozygote values at phenotypic levels *P*_*i*_ and *P*_*i*+1_ are respectively (see equation 1 in section 2.1 Theoretical Background):

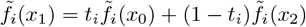

and

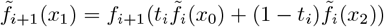

or, in terms of *t*_*i*+1_:

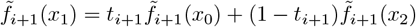

Using the properties of concave functions, we show from these expressions that *t*_*i*+1_ *< t*_*i*_, ∀*i* (**Appendix C**), therefore

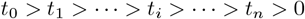

Non-additivity appears at the first phenotypic level. This effect then increases over successive levels (**Figure 2**B). The limit is *t*_∞_ = 0, which corresponds to the heterozygote value being identical to the value of individual *A*_2_*A*_2_, *i*.*e*. the high allele is completely dominant. An increase in dominance over phenotypic levels is the consequence of the increased curvature between the genotypic level and subsequent phenotypic levels (**Figure 2**A).

The previous results on phenotypic convergence and the increase in deviation from additivity are independent of the type of concave function. In **Appendix D**, we analytically derived cascades of hyperbolic functions and power functions and showed that results were quite similar.

#### Polygenic traits

For polygenic traits, the value of the L1*×*L2 hybrid can be expressed as a function of 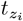, the deviation from additivity at level *i*:

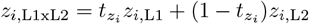

Combining this equation with equation 2, we obtain the following relationship between 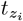 and the *t*_*i,j*_, the deviation from additivity of components *j*:

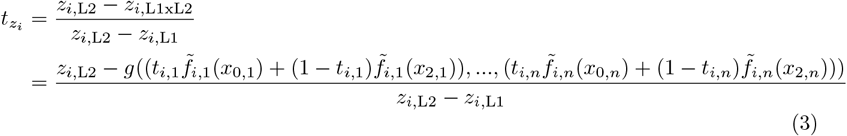

Unlike for monogenic traits, 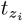 is not necessarily positive. The denominator *z*_*i*,L2_ *− z*_*i*,L1_ is positive, but the numerator *z*_*i*,L2_ *− z*_*i*,L1xL2_ will be negative if the hybrid value *z*_*i*,L1xL2_ is higher than the L2 value *z*_*i*,L2_. This can occur if lines have a complementary assortment of loci with high and low alleles, as heterozygosity masks the effect of low alleles in each line. Thus, we can distinguish two situations: (i) the hybrid value is higher than the mid-parental value but lower than the best parental value: 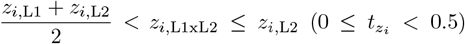. In genetic terms, this corresponds to mid-parent heterosis (MPH); (ii) the hybrid value is higher than the best-parental value: *z*_*i*,L1xL2_ > *z*_*i*,L2_ (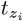 is therefore negative). This corresponds to best-parent heterosis (BPH). So, the range of variation of the deviation from additivity for polygenic traits is:

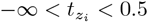

To put it in a less abstract way, let us illustrate this result using the example a digenic trait and two different parental line pairs. In one pair, line L2 has only high alleles (genotype *A*_2_*A*_2_*B*_2_*B*_2_) and line L1 has only low alleles (genotype *A*_1_*A*_1_*B*_1_*B*_1_) (**Figure 3**A, **3**C, **3**E and **3**G), while in the other both lines have a homozygous locus with low alleles (L2 has *A*_2_*A*_2_*B*_1_*B*_1_ and L1 has *A*_1_*A*_1_*B*_2_*B*_2_) (**Figures 3**B, **3**D, **3**F and **3**H). In the former case the hybrid value is equal to the mid-parental value only if both the function *g* and the functions 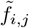 are linear (**Figure 3**A); otherwise there is MPH, *i*.*e*. 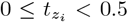 (**Figure 3**C, **3**E and **3**G). In the latter, the hybrid value is higher than the L2 value, and thus 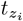 is negative (**Figure 3**F, **3**D and **3**H), except when functions *g* and 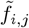 are linear (**Figure 3**B).

**Figure 3.**
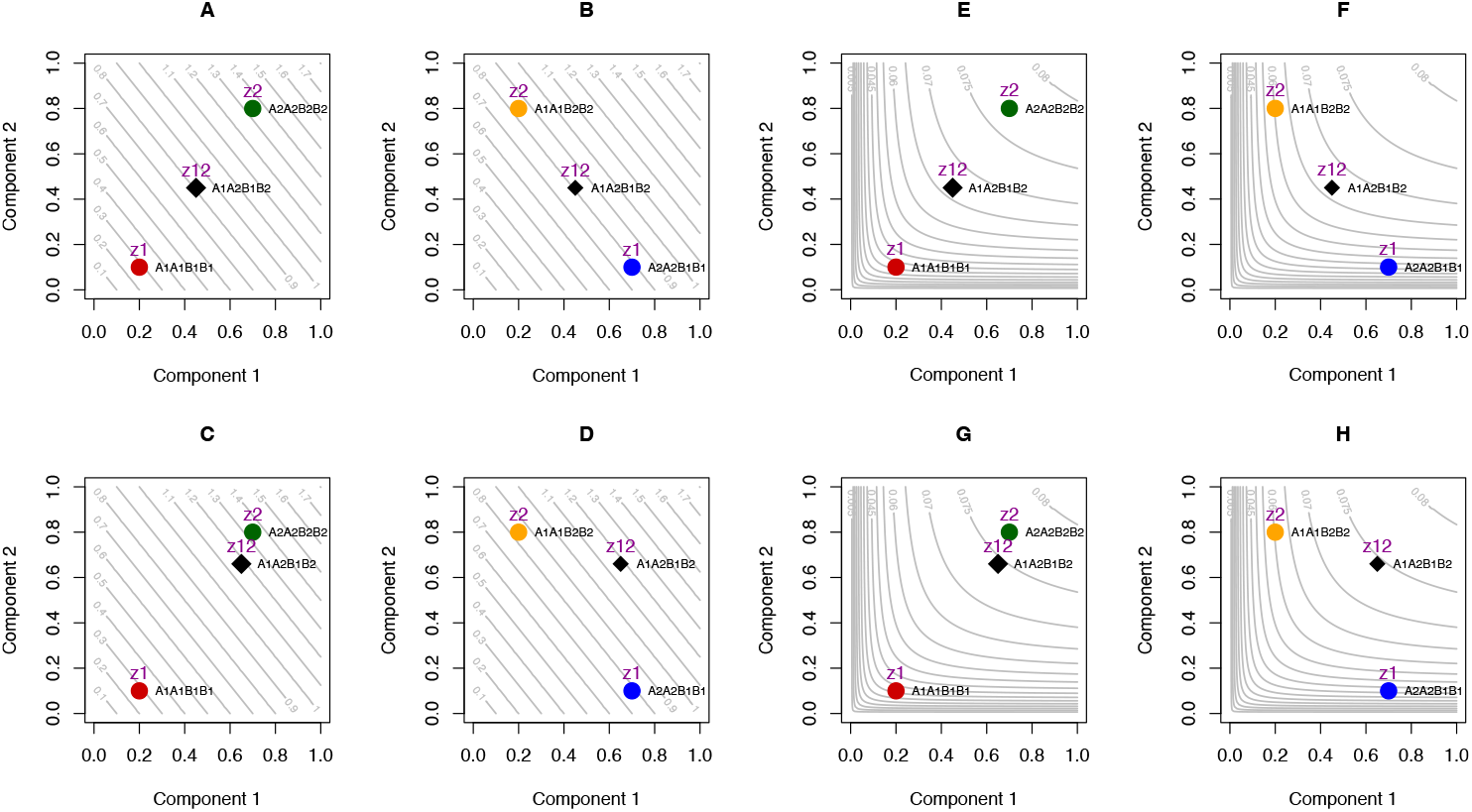
Deviation from additivity for a digenic trait. The trait value *z* (level curves) is related to the values of the two monogenic components 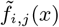 (axes) by the function 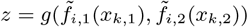 (see text). Function *g* is linear in **A, B, C** and **D** and concave in **E, F, G** and **H**. Functions 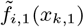 and 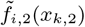 are linear in **A, B, E** and **F** (top row), and concave in **C, D, G** and **H** (bottom row). In **A, C, E** and **G** the cross is *A*_1_*A*_1_*B*_1_*B*_1_ *× A*_2_*A*_2_*B*_2_*B*_2_ (red and green dots, respectively). In **B, F, D** and **H**, the cross is *A*_2_*A*_2_*B*_1_*B*_1_ *× A*_1_*A*_1_*B*_2_*B*_2_ (blue and orange dots, respectively). Hybrids are represented by black diamonds. The highest hybrid value *z*_*i*,L1xL2_ relative to the line values *z*_*i*,L1_ and *z*_*i*,L2_ is observed in **G** and **H**, due to the concavity of both function *g* and functions 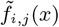, and to the genotypic complementarity of the lines. The deviation from additivity 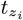 is negative, which corresponds to best-parent heterosis.

The extent of MPH or BPH depends on both the degree of concavity of function *g* and the deviation from additivity *t*_*i,j*_ of the components. Everything else being equal, high-level polygenic traits are expected to display more heterosis than the low-level traits because *t*_*i,j*_ decreases across the levels.

### 2.4 Two variants of the basic model

#### 2.4.1 Introducing a cost/crowding factor

Consider a genetically variable component of the transcription/translation machinery. Over a certain abundance, this component can have a biosynthetic cost and/or causes molecular crowding that hampers the output of the system, *e*.*g*. the rate of a metabolic flux. This can be modeled by introducing a cost/crowding factor (Alon, 2019; Kemble et al., 2020) to the function linking genotype *G* to the first phenotypic level *P*_1_:

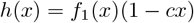

where *c* is the cost/crowding parameter, with 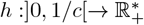. The function *h*(*x*) has a maximum, and the other relationships *P*_*i*_ ’ *P*_*i*+1_ remain ascending concave functions (**Figure 4A**).

**Figure 4.**
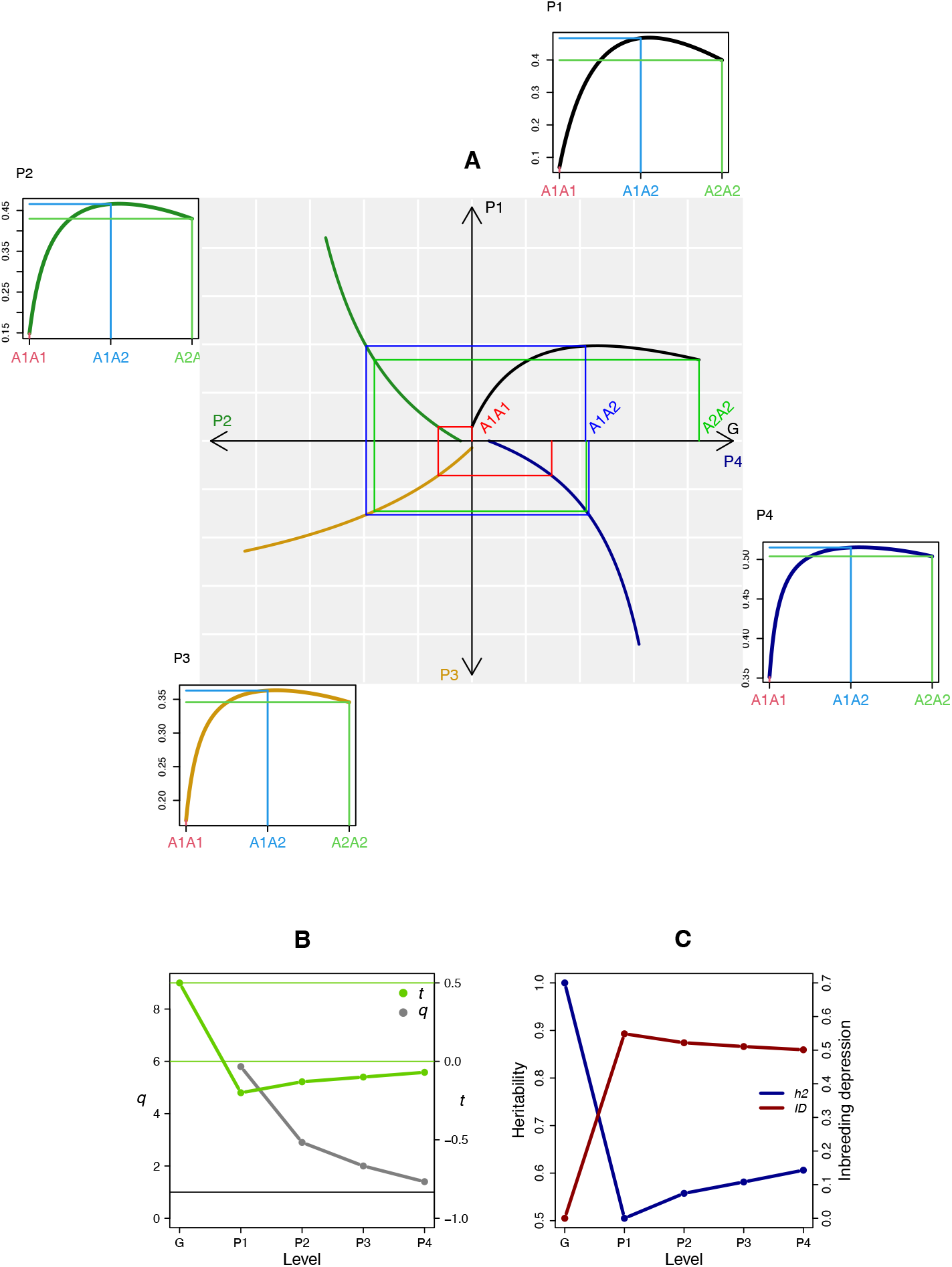
Phenotypic convergence and deviation from additivity across phenotypic levels when factoring in biosynthetic cost and/or crowding. The panels, symbols and colors are the same as in Figure 2. The functions for each level transition are the same, except for the *G → P*_1_ transition where the function includes a cost/crowding factor 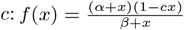, with *α* = 0.025, *β* = 0.36 and *c* = 0.47. **A** The effect of cost/crowding results in overdominance at level *P*_1_ (*t*_1_ *<* 0), but this effect diminishes from *P*_1_ to *P*_4_ (see the increase of *t*_*i*_ in **B**). Heritability and inbreeding depression vary accordingly (**C**).

Introducing a cost/crowding factor does not affect the convergence to 1 of the ratio *q*_*i*_, because the effect of cost/crowding does not alter the concavity of the functions (**Figure 4B**). If *h*(*x*_2_) *< h*(*x*_0_), *i*.*e*. if the effect of cost/crowding reverses the homozygote values between *G* and *P*_1_, the *q*_*i*_’s become lower than 1 but still converge to 1.

Regarding the deviation from additivity, we have the same expressions as in section 2.3:

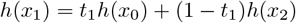

and

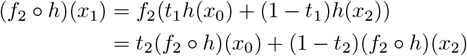

If *h*(*x*_2_) > *h*(*x*_1_), the inequality C.1 of **Appendix C** applies to *f*_2_ ∘ *h*:

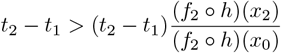

and we have as previously:

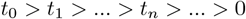

If *h*(*x*_2_) *< h*(*x*_1_), *i*.*e*. if there is overdominance, *t*_1_ is negative because *h*(*x*_1_) is over the upper limit of interval *{h*(*x*_0_), *h*(*x*_2_)*}*. Because *f*_2_ is an ascending concave function, (*f*_2_ ∘*h*)(*x*_1_) is also over the upper limit of the interval *{*(*f*_2_ ∘*h*)(*x*_0_), (*f*_2_ ∘*h*)(*x*_2_)*}*, therefore *t*_2_ is also negative, and so on for the subsequent *t*_*i*_. To determine the relationship order of the *t*_*i*_, we used the inequality *h*(*x*_0_) *< h*(*x*_2_) *< h*(*x*_1_), and showed that (**Appendix E**)

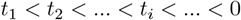

Thus, when the effect of cost/crowding results in overdominance, the successive *t*_*i*_ values converge to 0 as before, but by negative values (**Figure 4**B). In genetic terms, the overdominance progressively decreases, the limit for *t*_∞_ being complete dominance. Note that if a large cost/crowding effect makes *h*(*x*_2_) *< h*(*x*_0_), the conclusions are qualitatively the same, but the sign and the limit of *t*_*i*_ are modified: *t*_1_ is positive and higher than 1, then decreases toward 1 across the levels:

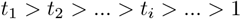

The limit 1 corresponds to complete dominance of the *A*_1_ allele over the *A*_2_ allele.

For polygenic traits, the effect of cost/crowding can increase the amplitude of heterosis because it increases the curvature of the hypersurface. However, this “excess” of heterosis will be less marked for the most integrated traits, in the same way that overdominance fades between the lowest and the highest levels, tending toward complete dominance.

#### 2.4.2 Phenotypic cascades with sigmoidal relationships

Transcription cascades (Lee et al., 2002; Manioudaki and Poirazi, 2013), protein kinase cascades (Ferrell Jr, 1997; Nijhout et al., 2003) and repressor protein-target gene expression cascades (Li et al., 2019) have sigmoidal relationships. As long as the genotypic variation is in the concave part of the curve, *i*.*e*. beyond the inflection point, previous conclusions apply. Otherwise, *q*_*i*_ can increase instead of decrease across phenotypic levels and negative non-additivity (*t* > 0.5) is observed. In **Appendix F**, we analyzed the variation of *t*_*i*_ and *q*_*i*_ across the phenotypic levels in two cascades involving such sigmoidal relationships: a three-step cascade from a transcription factor to growth and a cascade from three successive transcription factors to enzyme synthesis. In the latter, the steepness of the sigmoid increases as levels become more integrated, which may explain compensatory mutations and haploinsufficiency (see Bost and Veitia (2014) for a thorough analysis of a sigmoidal transcriptional activation cascade).

## 3 Implications for heritability and inbreeding depression

The increased concavity of the genotype-phenotype relationship across organization levels, which results in an increasing deviation from additivity, is expected to have two genetic consequences: a decrease of narrow-sense heritability and an increase in inbreeding depression.

### 3.1 Heritability decreases as the level of phenotypic integration increases

For monogenic biallelic variation, the additive and dominance variances under random mating are respectively (Falconer and Mackay, 1996)

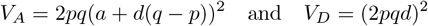

where *p* and *q* are the frequencies of the two alleles, *a* is the additive allelic effect and *d* is the dominance.

In a population derived from a cross between two inbred lines, we have *p* = *q* = 0.5, so these equations are reduced to:

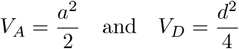

Thus, the narrow-sense heritability, 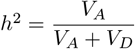 is:

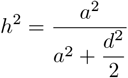

or, at any phenotypic level *i*:

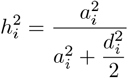

With our notations 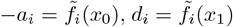 and 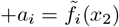. Since 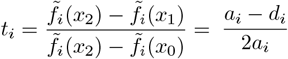, we have *d*_*i*_ */a*_*i*_ = 1 *−* 2*t*_*i*_, and:

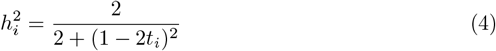

The first derivative of 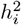 with respect to *t*_*i*_ is positive. As the most integrated traits have the lowest *t* values, *i*.*e*. the highest dominance, they are expected to display the lowest heritability values (**Figure 2**C, blue curve).

The decrease in heritability across the levels is also valid for polygenic traits, which are functions of monogenic components and are linked to the genotype along a concave hypersurface whose curvature is higher for the most integrated traits.

The effect of cost/crowding does not modify equation 4, and the variation of 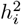 across the levels follows that of *t*_*i*_ (**Figure 4**B and **4**C). Heritability drops sharply at level *P*_1_, then increases slightly across higher levels, but remains much lower than in the absence of cost/crowding.

### 3.2 Inbreeding depression increases with the level of phenotypic integration

After *m* generations of heterozygote *A*_1_*A*_2_ self-fertilization in the absence of selection, the theoretical frequencies of genotypes *A*_1_ *A*_1_, *A*_1_*A*_2_ and *A*_2_*A*_2_ are respectively,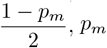 and 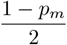, where 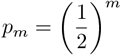. Thus, the mean phenotypic value of the population at level *P*_*i*_ is

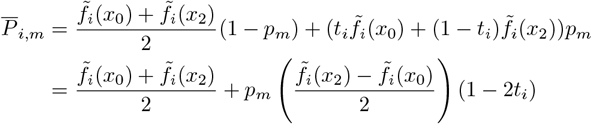

The inbreeding depression, *ID*_*i,m*_, can be quantified by:

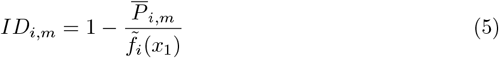

where 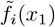 is the heterozygote value (Charlesworth and Willis, 2009). Since traits may have possibly different units of measure from one level to another, the difference 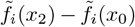 was set to 1 and 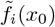 set to 0 to compare inbreeding depression across levels. Thus 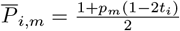 and 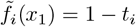, therefore

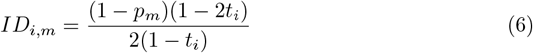

The first derivative of *ID*_*i,m*_ with respect to *t*_*i*_ is negative, so that inbreeding depression increases when *t* decreases, *i*.*e*. when phenotypic levels become more integrated (**Figure 2**C, red curve).

The same conclusions apply when considering polygenic traits, which are assumed to be *n*-dimensional concave functions of the components.

Factoring in cost/crowding does not modify equation 6, and we still have the inverse relationship between *t* and *ID* **(Figure 4**C). *ID* increases sharply at level *P*_1_, then decreases progressively, but remains higher than in absence of cost/crowding.

## 4 Results at a glance

In **Table 1**, we summarize the evolution of the phenotypic ratio *q*_*i*_ and the deviation from additivity *t*_*i*_ across phenotypic levels, as well as the genetic implications of an increase in deviation from additivity. Whether there is a constraint (cost/crowding) or not, there is both phenotypic convergence and an increase in dominance/heterosis as phenotypic levels become more integrated. As a consequence, heritability decreases and inbreeding depression increases.

**Table 1.** Genetic consequences of the propagation of genotypic variation across phenotypic levels when relationships between levels are concave. ^(*)^Or decreases to 1 if the effect of cost/crowding reverses the homozygote values.

| | Phenotypic ratio ($q_i$) | Deviation from additivity ($t_i$) |
| --- | --- | --- |
| Without<br>cost/crowding | *Decreases to 1*<br>→ Phenotypic convergence | *Decreases to 0*<br>→ Dominance and heterosis increase<br>→ Heritability decreases<br>→ Inbreeding depression increases |
| With<br>cost/crowding | *Decreases or increases to 1*<br>→ Phenotypic convergence | If $\tilde{f}_1(x_1) < \tilde{f}_1(x_2)$:<br>*Decreases to 0*<br>→ Same as above<br>If $\tilde{f}_1(x_1) > \tilde{f}_1(x_2)$:<br>*Increases to 0* (*)<br>→ Overdominance evolves<br>into complete dominance,<br>and strong heterosis gradually diminishes<br>→ Heritability drops markedly,<br>then increases only slightly<br>→ Inbreeding depression increases<br>markedly, then decreases only slightly |

## 5 Discussion

### 5.1 Genetic implications of the propagation of variation across phenotypic levels

At different levels of the biological hierarchy, diminishing returns and sigmoidal responses seem to be the most common types of relationship between phenotypic levels. As natural selection is assumed to keep genetic variation on the right-hand side of these curves because deleterious mutations are selected against (Kacser and Burns, 1981; Hartl et al., 1985; Tokuriki and Tawfik, 2009; Coton et al., 2022), it is relevant to use the theory of concave functions to describe the way genetic variation propagates across phenotypic levels.

We found that, as the phenotypic levels become more integrated: (i) phenotypic differences between individuals become less pronounced; (ii) the deviation from additivity – dominance for monogenic traits, heterosis for polygenic traits – increases (**Table 1**). These conclusions are valid regardless of the concave functions used, provided they are monotonically increasing.

The first result is fully consistent with the buffering of phenotypic variation observed from the transcriptome/proteome/metabolome levels up to the macroscopic levels. Indeed, most of the considerable variation observed at the low levels of biological organization in plants, animals and humans have no detectable cognate at the level of the macroscopic traits (Khaitovich et al., 2004; Fu et al., 2009; Connally et al., 2022). This multi-level degeneracy (Whitacre, 2012) may partly explain the pervasive robustness of biological systems (Félix and Barkoulas, 2015; Kitano, 2004), the phenotypic threshold effects (Rossignol et al., 2003), the developmental genetic basis of canalization (Hall-grimsson et al., 2019) and the selective neutrality of most molecular polymorphisms (Kimura, 1983; Ho et al., 2017; Zhang, 2018).

The second result – the increase in non-additivity across phenotypic levels – is consistent with the marked contrast between inheritance at the molecular level and at the macroscopic level. While dominance and heterosis are common and often extensive for integrated traits (*e*.*g*. Billiard et al., 2021; Flint-Garcia et al., 2009), this is not the case for molecular phenotypes such as protein, transcript and metabolite abundances. Additivity is generally prevalent for these low-level traits (Powell et al., 2013; Li et al., 2020; Zhou et al., 2019; Cui et al., 2006; Xing et al., 2016), and when there is nonadditivity, it is not at the same magnitude as for high-level traits. For instance Li et al. (2020) observed that in maize metabolic heterosis was relatively mild compared to the heterosis of integrated traits. Even between molecular traits there are differences that reflect their position in the intracellular hierarchy. For instance, Paschold et al. (2010) found five times more cases of heterosis for protein abundance than for gene expression in a maize hybrid, which is expected if we assume that protein abundance is a more integrated traits than transcript abundance.

High deviation from additivity for the most integrated traits may also explain the low heritability and high inbreeding depression of life history traits. Life history traits, or “fitness components”, are traits that affect fecundity, fertility, viability, survival, development rate, age and size at maturity, etc. Reviews in plants and animals (including humans) have repeatedly shown that fitness components generally have lower heritability than morphological, behavioral or physiological traits (Mousseau and Roff, 1987; Roff and Mousseau, 1987; Visscher et al., 2008; Yang, 2017). The reason traditionally given is that fitness components are more subject to natural selection than other traits that are less closely connected with fitness, and this reduces their additive genetic variance (Falconer and Mackay, 1996) (even though their additive variation coefficient may be quite high, as discussed in Flatt (2020)). Our results suggest another non-exclusive reason. In the phenotypic hierarchy, fitness components are considered to be high-level traits because they integrate various underlying structural, physiological and biochemical mechanisms (Arnold, 1983; Falconer and Mackay, 1996; Violle et al., 2007), and therefore their relationship with the genotype is assumed to be more concave than for lower-level traits. In genetic terms, this means high deviation from additivity, which leads to low heritability (**Figure 2**B and **2**C).

Regarding inbreeding depression, the literature shows that the fitness-related traits display more inbreeding depression upon selfing than less integrated traits (Good and Hallauer, 1977; Falconer and Mackay, 1996; DeRose and Roff, 1999). For instance the compilation by DeRose and Roff (1999) covering 86 life-history traits and 15 morphological traits in 54 animal species revealed that the former had a median reduction in trait value due to inbreeding (full-sib, *F* = 0.25) of ≈ 11.8%, whereas the latter had a reduction of ≈ 2.2%. The best explanation for this observation is to assume that dominance — and therefore heterosis for polygenic traits — is on average higher for fitness components than for other traits, due to the stronger curvature of the genotype-phenotype relationship, resulting in a positive relationship between inbreeding depression and phenotypic level.

### 5.2 The issue of cost and/or crowding

The prevalence of concave and plateauing genotype-phenotype relationships is congruent with the view of biological systems as physical/chemical networks of matter-energy exchanges. In such networks, the effect of increasing the efficiency of a single component is limited by the efficiency of all other components in the network, which results in diminishing returns. This has been formally proven in electrical circuits, which are to some extent analogous to metabolic networks (Petrizzelli et al., 2024). In biological networks, better component efficiency can also be achieved by increasing the component’s abundance, which is energetically costly and can exacerbate molecular crowding. It has been shown that molecular crowding can impose a physical limit on transcription and translation rates (Klumpp et al., 2013, 2019). Moreover, protein overproduction can lead to protein burden, which reduces cell growth (Kurland and Dong, 1996; Snoep et al., 1995; Eguchi et al., 2018; Kafri et al., 2016). Biosynthetic cost represents a selective constraint (Kafri et al., 2016) (Koehn, 1991; Vilaprinyo et al., 2010), which can lead to “competition” between enzymes for cell resources (Coton et al., 2022, 2023). An increase (resp. decrease) in the concentration of one or more enzyme due to genetic or environmental causes can result in a decrease (resp. increase) in the concentration of other enzymes (Snoep et al., 1995; Albertin et al., 2013).

If these phenomena were widespread, genotype-phenotype curves with a maximum should be commonly observed, which is not the case. There are two possible non-exclusive reasons for this. (i) The majority of the cell components are at low or very low concentration, so they contribute very little to the overall crowding and energy expenditure of the cell. Therefore, their overexpression has a negligible effect on the whole system. Note that protein burden effects have been experimentally detected using levels of gene overexpression that probably far exceed natural expression levels. (*e*.*g*. Snoep et al., 1995; Kurland and Dong, 1996; Eguchi et al., 2018); (ii) Existing concave genotype-phenotype curves with a maximum at the level of the transcription/translation machinery are buffered at subsequent phenotypic levels. To illustrate this point, we introduced an arbitrary cost/crowding factor to the function linking *G* to *P*_1_, the other *P*_*i*_ *→ P*_*i*+1_ relationships remaining ascending functions (**Figure 4**). Formally, this does not modify the results qualitatively: across levels, the ratio *q*_*i*_ between homozygote values at level *i* converges to 1 and the deviation from additivity *t*_*i*_ at level *i* converges to 0 from negative values. If the effect of cost/crowding reverses the order of homozygote values, then *q*_*i*_ is lower than 1 but still converges to 1, and *t*_*i*_ converges to 1 from values greater than 1. In both cases, this means that overdominance at the molecular level decreases across the higher levels, and tends toward complete dominance.

There are other molecular explanations for overdominance, such as the formation of favorable oligomeric proteins in heterozygotes and more balanced dosage effects in complex regulatory networks (reviewed in Liberatore et al., 2013; Fievet et al., 2018). However, the number of well-documented cases of overdominance at the macroscopic level is extremely small compared to the considerable number of genes showing complete or partial dominance. Furthermore, some cases of overdominance are actually due to antagonistic pleiotropy, in which two (or more) traits controlled by the same gene each exhibit dominance, but in opposite directions (ex. Allison, 1954; Dollinger, 1985; LaFountain et al., 2017). In any case, irrespective of the extent of overdominance at the molecular level, the phenotypic buffering effect from the lower levels to the more integrated levels is expected to gradually reduce overdominance until it can no longer be distinguished from complete dominance, which possibly explains why overdominance is so rare at macroscopic levels. On the other hand, heterosis for high-level traits is extremely common, not only in crops (Labroo et al., 2021) and livestock (Dickerson, 1973), but also in wild species, including microorganisms (*e*.*g*. Herbst et al., 2017). This is consistent with the fact that these traits: (i) are generally more polygenic than low-level traits, thus offering more possibilities for complementary genetic assortments; and (ii) have components with high dominance due to strong concavity at high levels of organization. From this perspective, heterosis can be defined as multidimensional dominance and appears to be an almost inevitable systemic effect.

### 5.3 Systemic *vs*. reductionist approaches

We are fully aware that this theoretical and geometric approach to the genotype-phenotype relationship ignores the extreme diversity and complexity of the molecular and cellular mechanisms that govern the relationships between phenotypic levels, as well as all the specificities of particular traits and functions. However, just because a system is incompletely described does not mean that generalizations are impossible, and to quote Dhar and Giuliani (2010) “the molecular level description is sometimes inadequate to explain higher-level behavior of organisms”. Our systemic approach leads to results that are fully consistent with various long-standing observations in quantitative and evolutionary genetics. The reason probably lies in the general and unavoidable constraint superimposed on all “local” mechanisms, namely the limitation matter and energy in the cell, which is likely to be the main explanation for the pervasiveness of concave curves with saturation.

In their article *Physics Behind Systems Biology*, Radde and Hütt (2016) stated that “physics has a long tradition of characterizing and understanding emergent collective behaviors in systems of interacting units and searching for universal laws”. This tradition is much less developed in biology, because biological processes are highly complex and diverse, and because natural variability tends to mask regularities. However, Mendel’s laws, the Hardy-Weinberg equilibrium and the principle of natural selection are examples of fruitful generalizations that play a central role in genetics and evolutionary biology. By working on a global rather than local scale, our study reveals general behaviors that shed light on the genetic and evolutionary implications of the genotype-phenotype relationship.

## Acknowledgements

We would like to warmly thank Dr. Catherine Damerval and Dr. Marianyela Petrizzelli for stimulating discussions and for their careful review of the manuscript. We thank Dr. Hélène Citerne for language corrections.

## 6 APPENDICES

### A List of symbols

| Symbol | Meaning |
| --- | --- |
| $G$ | Genotypic level |
| $P_i$ | Phenotypic level $i$ ($i \in [1, n]$) |
| $x_0, x_1, x_2$ | Genotypic values of homozygote $A_1A_1$, heterozygote $A_1A_2$ and homozygote $A_2A_2$, respectively |
| $f_i$ | Function relating $P_{i-1}$ and $P_i$ |
| $\tilde{f}_i$ | Function relating genotypic level $G$ and phenotypic level $P_i$ |
| $t_i$ | Deviation from additivity at level $i$ for function $\tilde{f}_i$ |
| $g$ | $n$-dimensional concave function of $n$ components |
| $z$ | Value of function $g$ |
| $n$ | Number of components of function $g$ |
| $k$ | Individual index |
| $q_i$ | Ratio of the homozygote values at level $P_i$ |
| $t_{z_i}$ | Deviation from additivity for polygenic function $g$ at level $i$ |
| $h$ | Function combining $f_1$ and a cost/crowding factor $c$ |
| $p, q$ | Frequency of alleles $A_1$ and $A_2$, respectively |
| $h^2$ | Heritability |
| $ID$ | Inbreeding depression |
| $m$ | Number of generations of inbreeding |

### B Proof of convergence of phenotypic values

We have to compare 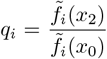 and 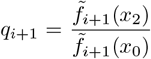, assuming that 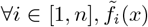 is an ascending concave function.

In **Figure B.1**, which shows an arbitrary relationship between 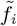 and 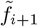, consider the straight line *y* = *ax* passing through the origin and intersecting the curve at point 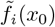. We have 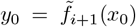. The slope of this line is 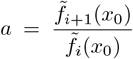, thus the ordinate of its point of intersection with the vertical line with abscissa 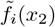 is 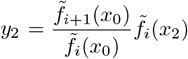. Thus

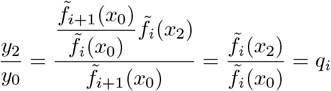

As the representative curve of a concave function is below each of its tangents, and *a* is greater than the slope of the tangent at point 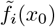, we have

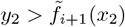

Thus

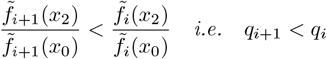

and by recurrence we have

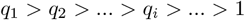

The limit is 1 because we assumed that these concave functions are all monotonically increasing.

**Figure B.1.**
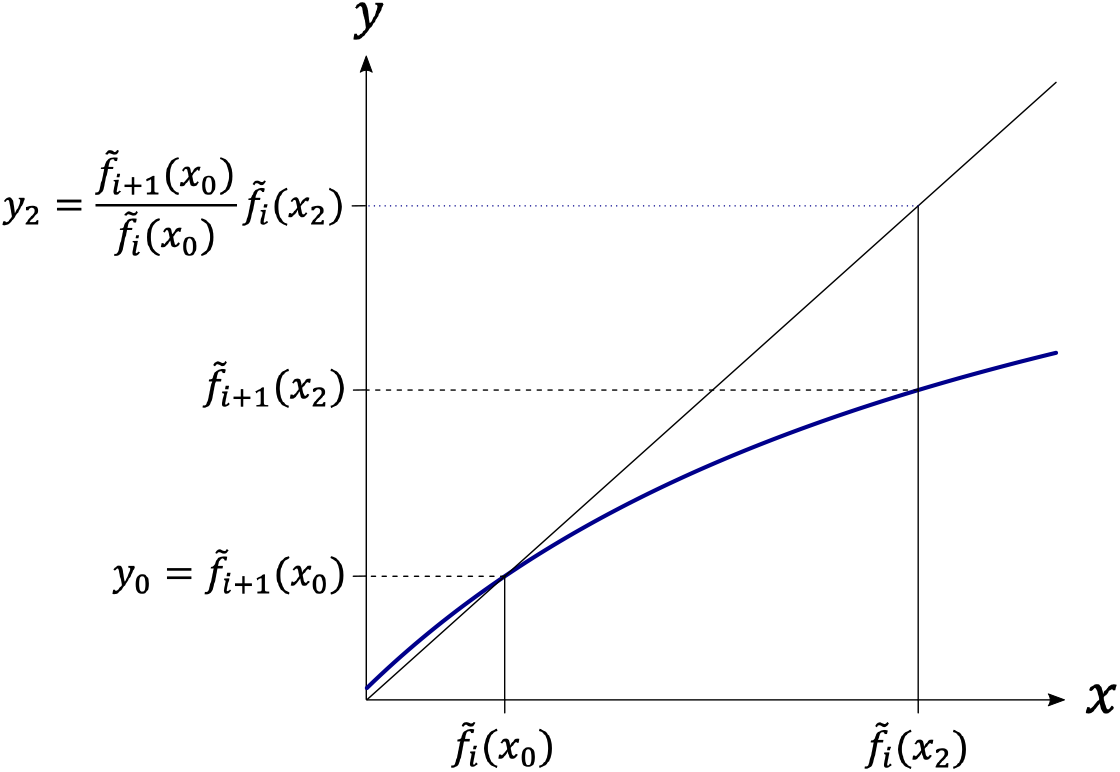
Relationship between the homozygote phenotypic values at levels *P*_*i*_ (*x*-axis) and *P*_*i*+1_ (*y*-axis).

### C Proof of increased deviation from additivity

The heterozygote value at level *P*_*i*_ is (equation 1 in the main text):

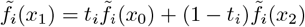

so at level *P*_*i*+1_:

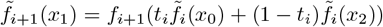

In terms of *t*_*i*+1_, we have:

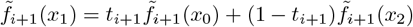

By the standard concavity argument applied to 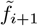, we have from the equality 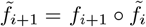:

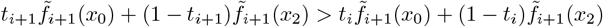

or

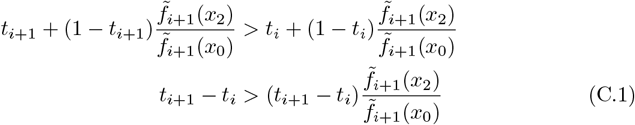

Because *f*_*i*_ is strictly increasing 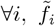 is also increasing ∀*i*. So 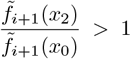 and inequality C.1 is respected only if *t*_*i*+1_ *− t*_*i*_ *<* 0, *i*.*e. t*_*i*+1_ *< t*_*i*_. We have *t*_0_ = 0.5 > *t*_1_ and by recurrence, we get *t*_*i*+1_ *< t*_*i*_ ∀*i*, thus

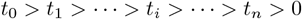

### D Two examples of cascades of concave functions

#### D.1 Hyperbolic functions

We consider a cascade of identical hyperbolic functions of the form *f* (*x*) = *x/*(1 + *x*).

The values of the successive levels are:

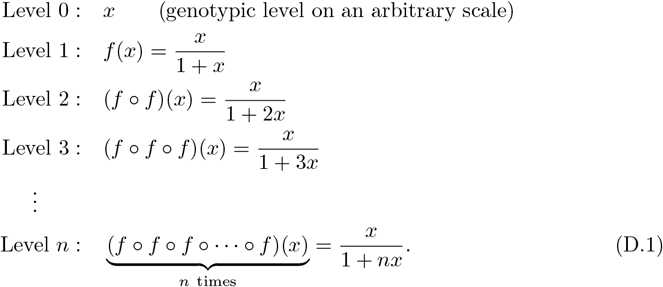

At the *n*^th^ level, the ratio of parental values is:

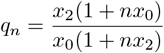

and the deviation from additivity is:

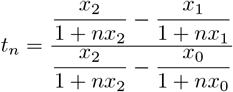

**Figures D.1**A, **D.1**B and **D.1**C show the increase of the curvature of the relationship between *x* and (*f* ∘ *f* ∘ · · · ∘ *f*)(*x*), the convergence of the phenotypic values of the parents (decrease of *q*_*n*_) and the increase of the deviation from additivity (decrease of *t*_*n*_) as phenotypic levels become more integrated.

#### D.2 Power functions

We consider a cascade of identical power functions of the form *f* (*x*) = *x*^*a*^, where *a* is a positive constant.

The values of the successive levels are:

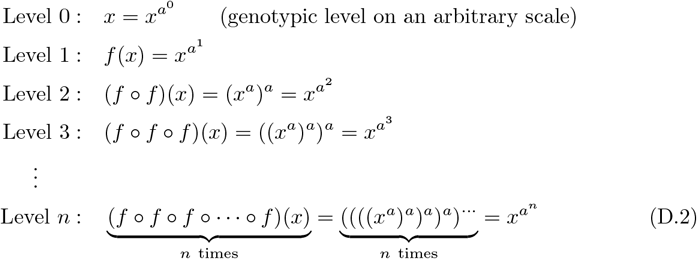

At the *n*^th^ level, the ratio *q*_*n*_ of the parental values is:

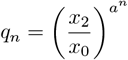

and the deviation from additivity is:

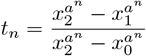

**Figures D.1**D, **D.1**E and **D.1**F show the increase of the concavity of the relationship between *x* and (*f* ∘ *f* ∘· · · ∘ *f*)(*x*), the convergence of the phenotypic values of the parents (decrease of *q*_*n*_) and the increase of the deviation from additivity (decrease of *t*_*n*_) as phenotypic levels become more integrated.

**Figure D.1.**
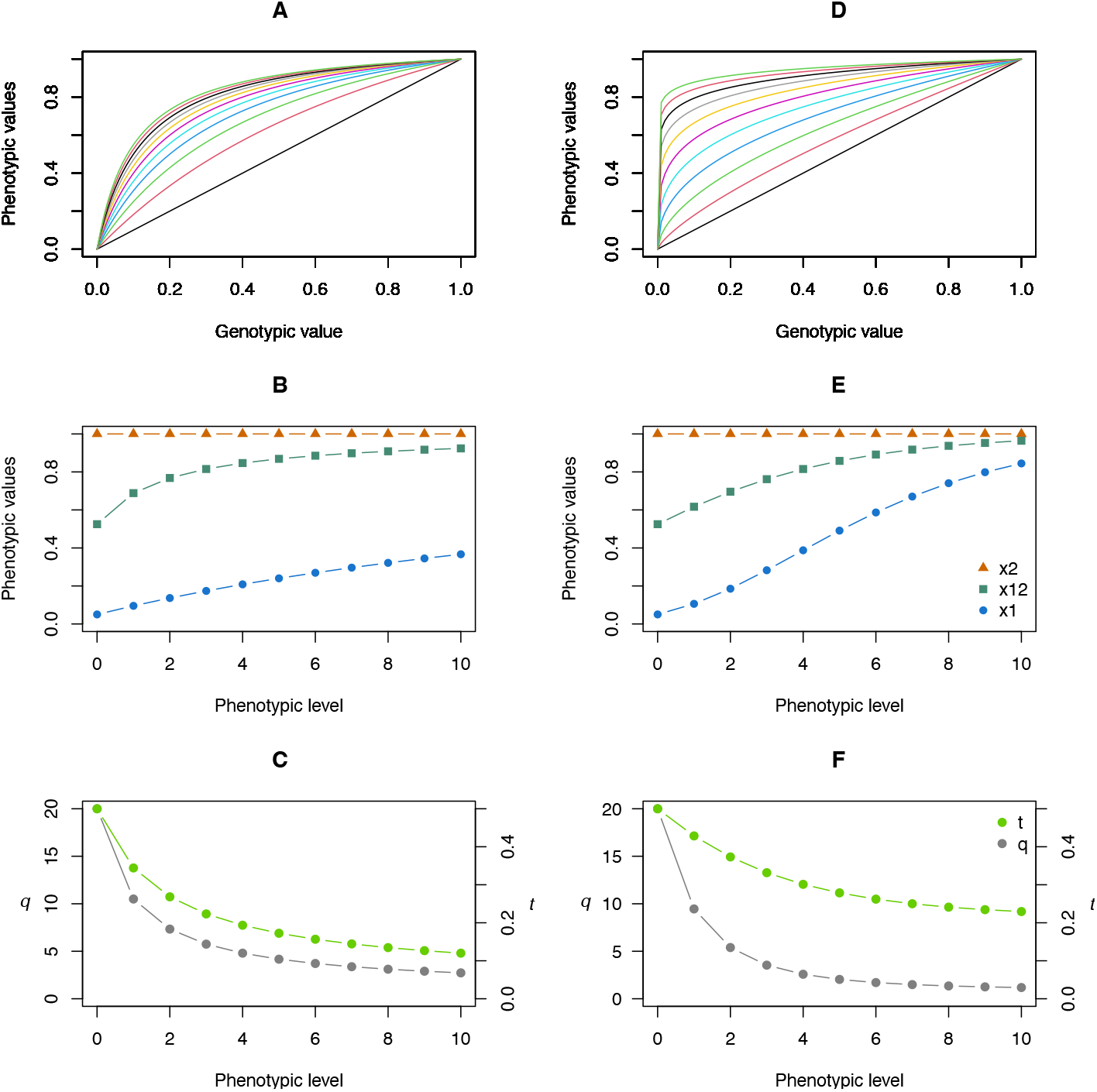
Phenotypic variation across 10 levels. A, B and C: hyperbolic relationships (normalized values). The function is 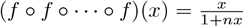, with *n* ∈ [0, 10] corresponding the level number. A: relationship between the genotypic value and the phenotypic values of the successive levels. The black straight line corresponds to *f* (*x*) = *x* (*n* = 0). B: variation of the phenotypic values across levels for the parental genotypic values *x*_0_ = 0.05 (blue dots), *x*_2_ = 1 (orange triangles) and that of their hybrid 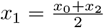 (green squares). C: ratio of parental values *q* (gray dots, scale given on the vertical axis on the left) and deviation from additivity *t* (green dots, scale given on the vertical axis on the right) decrease as the phenotypic levels increase. D, E and F: Power functions of the form 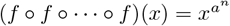, with *a* = 0.75 and *n* ∈ [0, 10]. Same representations as in A, B and C.

### E Deviation from additivity in the case of overdominance at the first phenotypic level

If the effect of cost/crowding results in overdominance at phenotypic level *P*_1_, the deviation from additivity *t*_1_ is negative. To determine the relationship order of the *t*_*i*_, we used the inequality *h*(*x*_0_) *< h*(*x*_2_) *< h*(*x*_1_), which makes it possible to define *τ*_1_ and *τ*_2_ such that

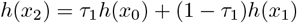

and

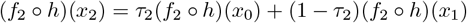

Due to the concavity of *h* and *f*_2_ ∘*h*, we get (using the same approach as previously, see inequality C.1)

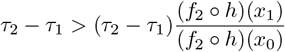

which is satisfied only if *τ*_2_ *< τ*_1_ because (*f*_2_ ∘ *h*)(*x*_1_) > (*f*_2_ ∘ *h*)(*x*_0_). The relationship order between *t*_2_ and *t*_1_ can be derived from the relationship *τ*_2_ *< τ*_1_:

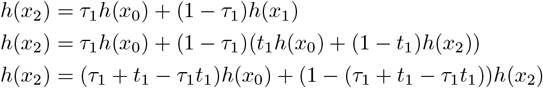

The last equality is met if *τ*_1_ + *t*_1_ *− τ*_1_*t*_1_ = 0, *i*.*e*. if 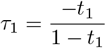. Similarly, 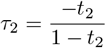. Since *τ*_2_ *< τ*_1_, we have 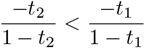, or 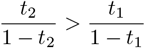 because *t <* 0 and 1*−t* > 0. Therefore, *t*_2_(1 *− t*_1_) > *t*_1_(1 *− t*_2_), thus *t*_2_ > *t*_1_, and by recurrence:

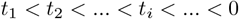

### F Cascades of sigmoid and concave functions

#### F.1 From transcription factor abundance to growth rate

We will consider three successive functions relating four phenotypic levels:

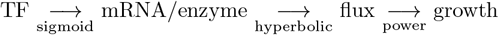

The sigmoid function relating a TF concentration and a gene expression level (or an mRNA concentration) is assumed to be a Hill function:

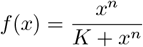

where *n* and *K* are constants.

The hyperbolic function relating enzyme concentration and a metabolic flux, assuming a linear relation between mRNA and enzyme concentrations, is assumed to be:

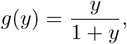

The function relating this flux and a macroscopic trait (*i*.*e*. the growth rate) is assumed to be a power function:

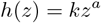

Therefore, the successive phenotypic values are given by the functions:

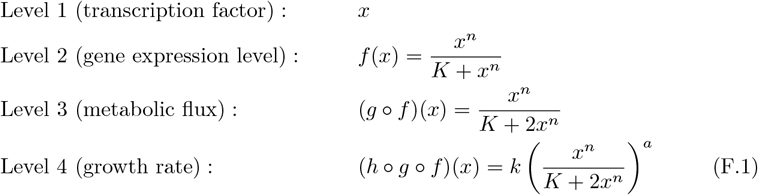

As shown **Figure F.1**A, the concavity of the right part of the curves increases with the phenotypic levels, while the convexity generated by the cooperative interactions between TF and DNA binding sites decreases, as can be seen by the decrease in the value of the inflection points. Consequently, the ratio of parental values *q* and the inheritance *t* will depend both on the initial parental concentrations *x*_1_ and *x*_2_ of the TF and on the phenotypic level considered. For instance, if *x*_1_ and *x*_2_ are on either side of the inflection point of the Hill function, ratio *q* for the mRNA level is higher than for TF, then decreases for growth rate but remains higher than the initial value. The deviation from additivity *t* exceeds 0.5 for the mRNA level then decreases below 0.5 for the growth rate (**Figures F.1**B and **F.1**D). With higher *x*_1_ and *x*_2_ values, *q* and *t* variation profiles of variation approach those of fully concave functions (**Figures F.1**C and **F.1**F).

#### F.2 Cascade of transcription factors upstream of enzyme synthesis

We will consider a cascade of three transcription factors (TF):

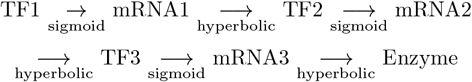

The relationships TF *→* mRNA are sigmoidal of the same form as previously:

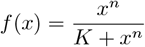

and the relationships mRNA *→* TF (or enzyme) are concave of the form:

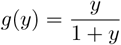

**Figure F.1.**
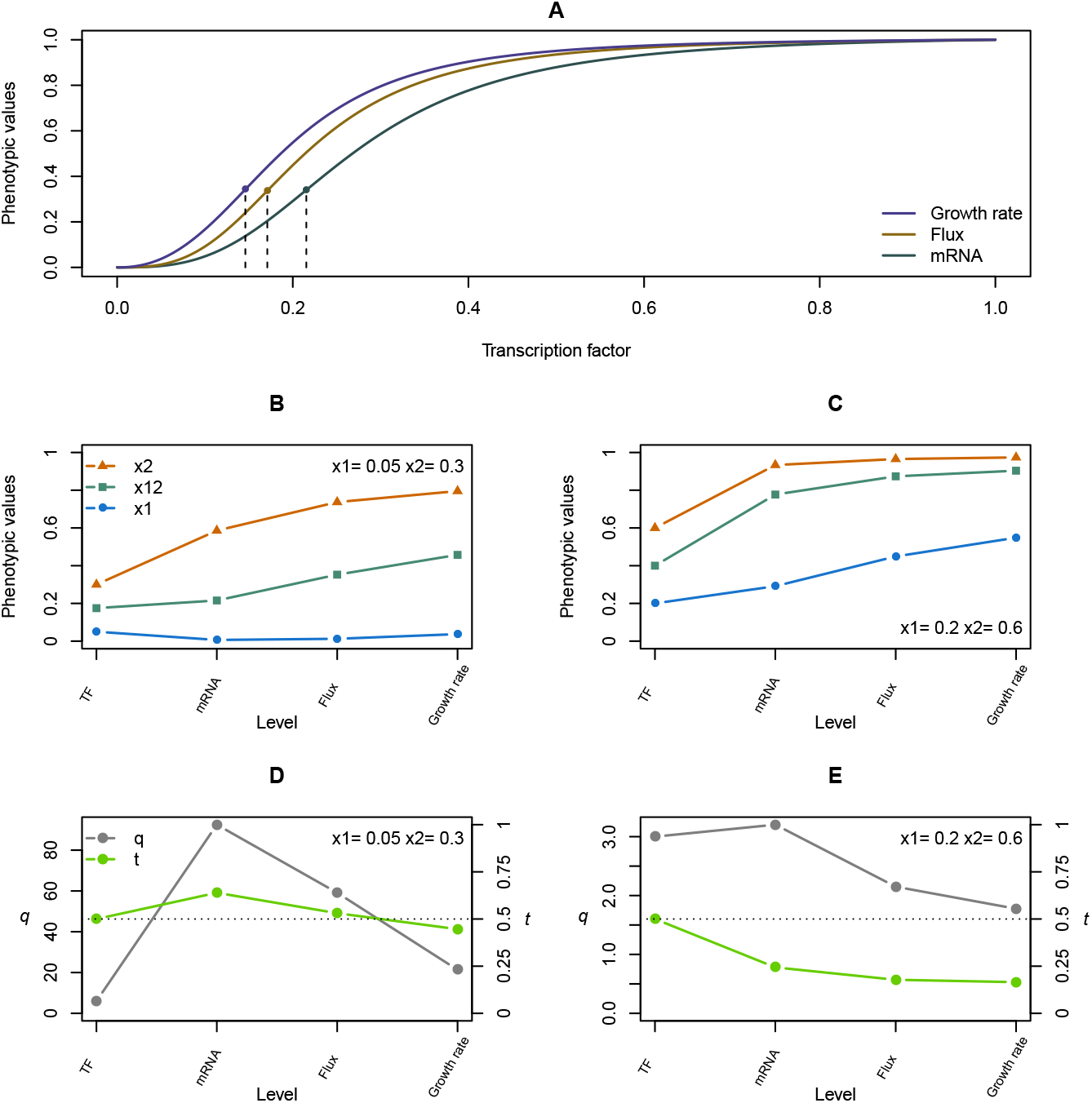
Sequence of sigmoid, hyperbolic and power functions. A. Relationship between the concentration of a transcription factor and three more integrated traits: mRNA abundance, metabolic flux and growth rate. Dotted lines indicate the inflection points of each curve. The parameters of the Hill function are *n* = 3 and *K* = 0.02, and of the power function *k* = 1 and *a* = 0.75. B and C. Variation of phenotypic values across the phenotypic levels for two pairs of parental values and their hybrids (resp. *x*_1_, *x*_2_ and *x*_12_). D and E. Variation of the ratio of the parental values, *q*, and of the deviation from additivity, *t*, across the phenotypic levels. Phenotypic values are normalized.

Therefore, the successive phenotypic values are given by the functions:

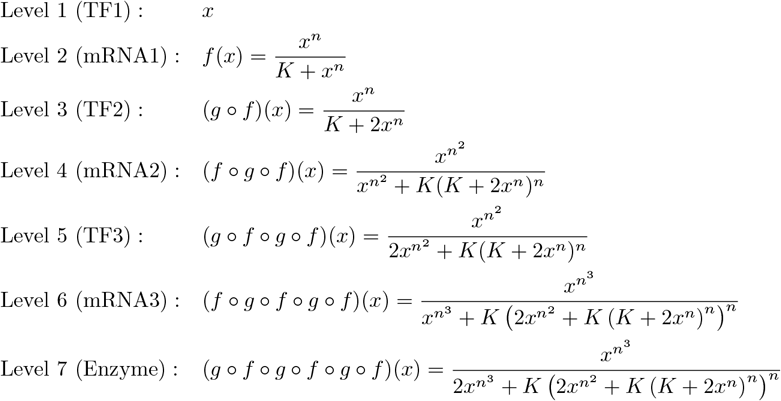

Due to the alternation of sigmoid and hyperbolic functions, the variation in curvature is no longer monotonous across levels, however there is a clear increase in the steepness of the sigmoid which becomes close to a step function at the enzyme level (**Figure F.2**A). The phenotypic variation across the levels and the *q* and *t* values, are very sensitive to the initial TF parental values (**Figures F.2**B to **F.2**E). Moreover if the initial TF values are both either below or above the inflection point of the Hill function, the question of the inheritance at the enzyme level is almost pointless since the parental values are very close.

**Figure F.2.**
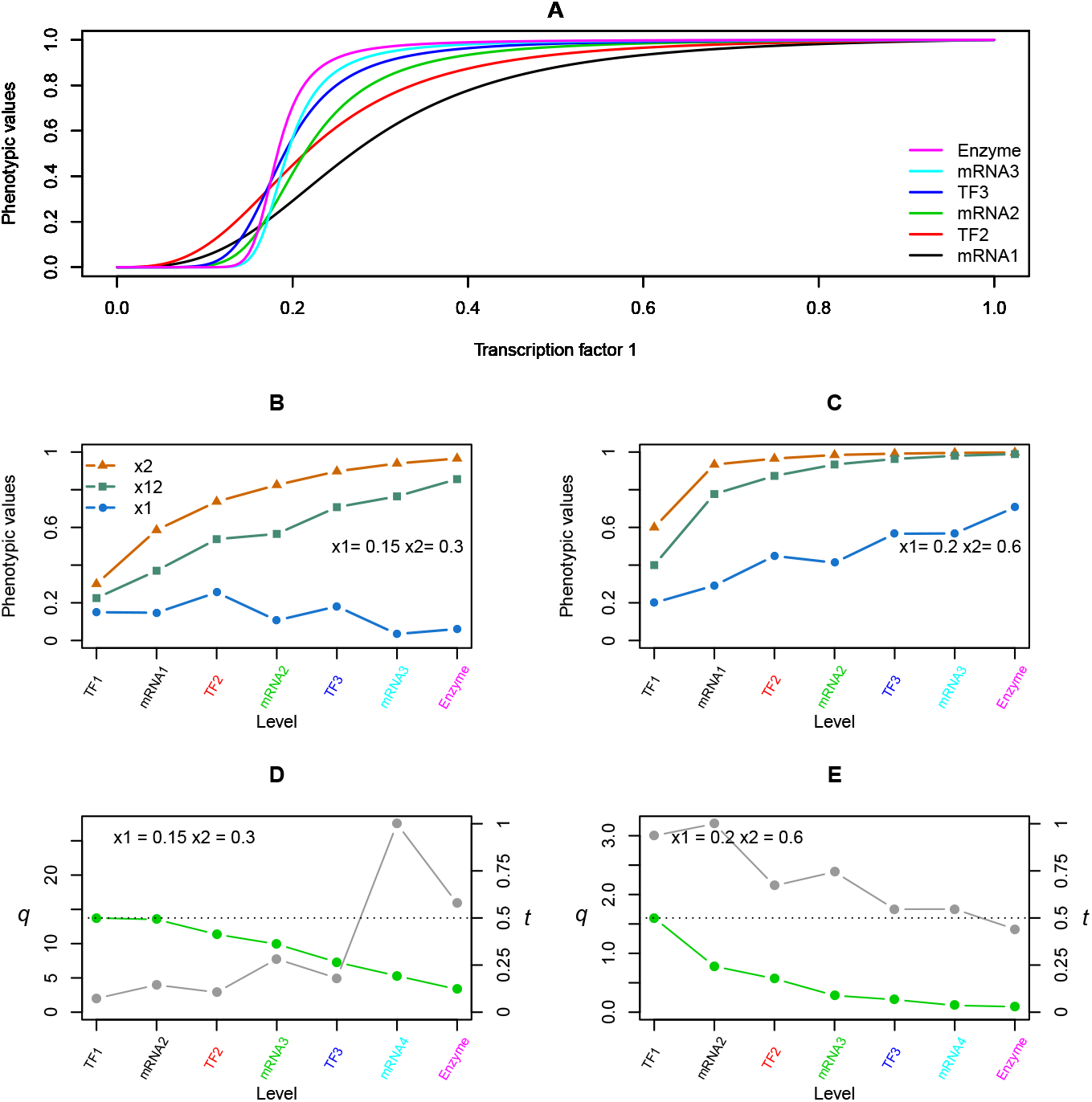
Cascade of three transcription factors. **A**. Relationship between the concentration of a transcription factor, TF1, and six successive phenotypic traits: mRNA1, TF2, mRNA2, TF3, mRNA3 and Enzyme. The parameters of the Hill function are *n* = 3 and *K* = 0.02. **B** and **C**. Variation of phenotypic values across the phenotypic levels for two pairs of parental values and their hybrids (resp. *x*_1_, *x*_2_ and *x*_12_). **D** and **E**. Variation of the ratio of the parental values, *q*, and of the deviation from additivity, *t*, across the phenotypic levels. Phenotypic values are normalized.

## Notes

### Competing Interest Statement

The authors have declared no competing interest.

## References

Albertin, W., Marullo, P., Bely, M., Aigle, M., Bourgais, A., Langella, O., Balliau, T., Chevret, D., Valot, B., Silva, T. d., Dillmann, C., Vienne, D. d., and Sicard, D. (2013). Linking Post-Translational Modifications and Variation of Phenotypic Traits. Molecular & Cellular Proteomics, 12(3):720–735.

Allison, A. (1954). Protection Afforded by Sickle-Cell Trait Against Subtertian Malarial Infection. Br. Med. J., 1(4857):290–294.

Alon, U. (2019). An Introduction to Systems Biology: Design Principles of Biological Circuits. Chapman & Hall.

Arnold, S. J. (1983). Morphology, Performance and Fitness. American Zoologist, 23(2):347–361.

Billiard, S., Castric, V., and Llaurens, V. (2021). The integrative biology of genetic dominance. Biological Reviews, 96(6):2925–2942.

Bost, B. and Veitia, R. A. (2014). Dominance and interloci interactions in transcriptional activation cascades: Models explaining compensatory mutations and inheritance patterns. Bioessays, 36(1):84–92.

Bunge, M. (1977). Levels and reduction. American Journal of Physiology-Regulatory, Integrative and Comparative Physiology, 233(3):R75–R82.

Campbell, D. T. (1974). ‘Downward Causation’ in Hierarchically Organised Biological Systems. In Ayala, F. J.and Dobzhansky, T., editors, Studies in the Philosophy of Biology: Reduction and Related Problems, pages 179–186. Macmillan Education UK, London.

Carey, M. (1998). The enhanceosome and transcriptional synergy. Cell, 92(1):5–8.

Charlesworth, D. and Willis, J. H. (2009). The genetics of inbreeding depression. Nat. Rev. Genet., 10(11):783–796.

Connally, N. J., Nazeen, S., Lee, D., Shi, H., Stamatoyannopoulos, J., Chun, S., Cotsapas, C., Cassa, C. A., and Sunyaev, S. R. (2022). The missing link between genetic association and regulatory function. Elife, 11:e74970.

Coton, C., Dillmann, C., and de Vienne, D. (2023). Evolution of enzyme levels in metabolic pathways: A theoretical approach. Part 2. Journal of Theoretical Biology, 558:111354.

Coton, C., Talbot, G., Le Louarn, M., Dillmann, C., and de Vienne, D. (2022). Evolution of enzyme levels in metabolic pathways: A theoretical approach. Part 1. Journal of Theoretical Biology, 538:111015.

Crow, J. F. and Kimura, M. (1970). An introduction to population genetics theory. An introduction to population genetics theory.

Cui, X., Affourtit, J., Shockley, K. R., Woo, Y., and Churchill, G. A. (2006). Inheritance Patterns of Transcript Levels in F1 Hybrid Mice. Genetics, 174(2):627–637.

de Vienne, D. (2022). What is a phenotype? History and new developments of the concept. Genetica, 150(3-4):153–158.

de Vienne, D., Coton, C., and Dillmann, C. (2023). The genotype–phenotype relationship and evolutionary genetics in the light of the Metabolic Control Analysis. Biosystems, 232:105000.

DeRose, M. A. and Roff, D. A. (1999). A comparison of inbreeding depression in lifehistory and morphological traits in animals. Evolution, 53(4):1288–1292.

Dhar, P. K. and Giuliani, A. (2010). Laws of biology: why so few? Syst Synth Biol, 4(1):7–13.

Dickerson, G. E. (1973). Inbreeding and heterosis in animals. Journal of animal science, 1973(Symposium):54–77.

Dollinger, E. J. (1985). Effects of Visible Recessive Alleles on Vigor Characteristics in a Maize Hybrid. Crop Science, 25(5):819–821.

Domingo, J., Baeza-Centurion, P., and Lehner, B. (2019). The Causes and Consequences of Genetic Interactions (Epistasis). Annu. Rev. Genom. Hum. Genet., 20(1):433–460.

Eguchi, Y., Makanae, K., Hasunuma, T., Ishibashi, Y., Kito, K., and Moriya, H. (2018). Estimating the protein burden limit of yeast cells by measuring the expression limits of glycolytic proteins. Elife, 7:e34595. Publisher: eLife Sciences Publications, Ltd.

Eldredge, N., Pievani, T., Serrelli, E., and Tëmkin, I. (2019). Evolutionary theory: a hierarchical perspective. University of Chicago Press.

Falconer, D. S. and Mackay, T. F. C. (1996). Introduction to quantitative genetics. London, Prentice Hall.

Ferrell Jr, J. E. (1997). How responses get more switch-like as you move down a protein kinase cascade. Trends in biochemical sciences, 22(8):288–289.

Fievet, J. B., Nidelet, T., Dillmann, C., and de Vienne, D. (2018). Heterosis Is a Systemic Property Emerging From Non-linear Genotype-Phenotype Relationships: Evidence From in Vitro Genetics and Computer Simulations. Front. Genet., 9:159.

Firczuk, H., Kannambath, S., Pahle, J., Claydon, A., Beynon, R., Duncan, J., Westerhoff, H., Mendes, P., and McCarthy, J. E. (2013). An in vivo control map for the eukaryotic mRNA translation machinery. Molecular systems biology, 9(1):635.

Flatt, T. (2020). Life-history evolution and the genetics of fitness components in Drosophila melanogaster. Genetics, 214(1):3–48.

Flint-Garcia, S. A., Buckler, E. S., Tiffin, P., Ersoz, E., and Springer, N. M. (2009). Heterosis Is Prevalent for Multiple Traits in Diverse Maize Germplasm. Plos One, 4(10):e7433.

Fu, J., Keurentjes, J. J., Bouwmeester, H., America, T., Verstappen, F. W., Ward, J. L., Beale, M. H., De Vos, R. C., Dijkstra, M., and Scheltema, R. A. (2009). Systemwide molecular evidence for phenotypic buffering in Arabidopsis. Nature genetics, 41(2):166–167.

Félix, M.-A. and Barkoulas, M. (2015). Pervasive robustness in biological systems. Nat. Rev. Genet., 16(8):483–496.

Gibson, G. (1996). Epistasis and pleiotropy as natural properties of transcriptional regulation. Theoretical population biology, 49(1):58–89.

Giorgetti, L., Siggers, T., Tiana, G., Caprara, G., Notarbartolo, S., Corona, T., Pasparakis, M., Milani, P., Bulyk, M. L., and Natoli, G. (2010). Noncooperative Interactions between Transcription Factors and Clustered DNA Binding Sites Enable Graded Transcriptional Responses to Environmental Inputs. Molecular Cell, 37(3):418–428.

Good, R. and Hallauer, A. R. (1977). Inbreeding Depression in Maize by Selfing and Full-sibbing. Crop Science, 17(6):935–940.

Hallgrimsson, B., Green, R. M., Katz, D. C., Fish, J. L., Bernier, F. P., Roseman, C. C., Young, N. M., Cheverud, J. M., and Marcucio, R. S. (2019). The developmental-genetics of canalization. In Seminars in cell & developmental biology, volume 88, pages 67–79. Elsevier.

Hartl, D., Dykhuizen, D., and Dean, A. (1985). Limits of Adaptation the Evolution of Selective Neutrality. Genetics, 111(3):655–674.

Herbst, R. H., Bar-Zvi, D., Reikhav, S., Soifer, I., Breker, M., Jona, G., Shimoni, E., Schuldiner, M., Levy, A. A., and Barkai, N. (2017). Heterosis as a consequence of regulatory incompatibility. BMC Biol, 15(1):38.

Ho, W.-C., Ohya, Y., and Zhang, J. (2017). Testing the neutral hypothesis of phenotypic evolution. Proc. Natl. Acad. Sci. U.S.A., 114(46):12219–12224.

Kacser, H. and Burns, J. (1981). The Molecular Basis of Dominance. Genetics, 97(3-4):639–666.

Kafri, M., Metzl-Raz, E., Jona, G., and Barkai, N. (2016). The cost of protein production. Cell reports, 14(1):22–31.

Kauffman, S. A. (1993). The origins of order: Self-organization and selection in evolution. Oxford University Press, USA.

Kazemian, M., Pham, H., Wolfe, S. A., Brodsky, M. H., and Sinha, S. (2013). Widespread evidence of cooperative DNA binding by transcription factors in Drosophila development. Nucleic Acids Research, 41(17):8237–8252.

Kemble, H., Eisenhauer, C., Couce, A., Chapron, A., Magnan, M., Gautier, G., Le Nagard, H., Nghe, P., and Tenaillon, O. (2020). Flux, toxicity, and expression costs generate complex genetic interactions in a metabolic pathway. Science Advances, 6(23):eabb2236.

Kemble, H., Nghe, P., and Tenaillon, O. (2019). Recent insights into the genotype–phenotype relationship from massively parallel genetic assays. Evol. Appl, 12(9):1721–1742.

Keren, L., Hausser, J., Lotan-Pompan, M., Slutskin, I. V., Alisar, H., Kaminski, S., Weinberger, A., Alon, U., Milo, R., and Segal, E. (2016). Massively parallel interrogation of the effects of gene expression levels on fitness. Cell, 166(5):1282–1294.

Khaitovich, P., Weiss, G., Lachmann, M., Hellmann, I., Enard, W., Muetzel, B., Wirkner, U., Ansorge, W., and Pääbo, S. (2004). A neutral model of transcriptome evolution. PLoS biology, 2(5):e132.

Kimura, M. (1983). The neutral theory of molecular evolution. Cambridge University Press.

Kitano, H. (2004). Biological robustness. Nature Reviews Genetics, 5(11):826–837.

Klumpp, S., Bode, W., and Puri, P. (2019). Life in crowded conditions: Molecular crowding and beyond. The European Physical Journal Special Topics, 227:2315–2328.

Klumpp, S., Scott, M., Pedersen, S., and Hwa, T. (2013). Molecular crowding limits translation and cell growth. PNAS, 110(42):16754–16759.

Koehn, R. K. (1991). The cost of enzyme synthesis in the genetics of energy balance and physiological performance. Biological Journal of the Linnean Society, 44(3):231–247.

Korn, R. W. (2005). The Emergence Principle in Biological Hierarchies. Biol Philos, 20(1):137–151.

Kurland, C. G. and Dong, H. (1996). Bacterial growth inhibition by overproduction of protein. Molecular microbiology, 21(1):1–4.

Labroo, M. R., Studer, A. J., and Rutkoski, J. E. (2021). Heterosis and hybrid crop breeding: a multidisciplinary review. Frontiers in Genetics, 12:643761.

LaFountain, A. M., Chen, W., Sun, W., Chen, S., Frank, H. A., Ding, B., and Yuan, Y.-W. (2017). Molecular basis of overdominance at a flower color locus. G3: Genes, Genomes, Genetics, 7(12):3947–3954.

Lee, T. I., Rinaldi, N. J., Robert, F., Odom, D. T., Bar-Joseph, Z., Gerber, G. K., Hannett, N. M., Harbison, C. T., Thompson, C. M., Simon, I., Zeitlinger, J., Jennings, E. G., Murray, H. L., Gordon, D. B., Ren, B., Wyrick, J. J., Tagne, J.-B., Volkert, T. L., Fraenkel, E., Gifford, D. K., and Young, R. A. (2002). Transcriptional Regulatory Networks in Saccharomyces cerevisiae. Science, 298(5594):799–804.

Li, X., Lalić, J., Baeza-Centurion, P., Dhar, R., and Lehner, B. (2019). Changes in gene expression predictably shift and switch genetic interactions. Nature communications, 10(1):3886.

Li, Z., Zhu, A., Song, Q., Chen, H. Y., Harmon, F. G., and Chen, Z. J. (2020). Temporal regulation of the metabolome and proteome in photosynthetic and photorespiratory pathways contributes to maize heterosis. The Plant Cell, 32(12):3706–3722.

Liberatore, K. L., Jiang, K., Zamir, D., and Lippman, Z. B. (2013). Heterosis: The Case for Single-Gene Overdominance. In Chen, Z. J.and Birchler, J. A., editors, Polyploid and Hybrid Genomics, pages 137–152. Wiley, 1 edition.

MacLean, R. C. (2010). Predicting epistasis: an experimental test of metabolic control theory with bacterial transcription and translation. Journal of Evolutionary Biology, 23(3):488–493.

MacLean, R. C., Perron, G. G., and Gardner, A. (2010). Diminishing returns from beneficial mutations and pervasive epistasis shape the fitness landscape for rifampicin resistance in Pseudomonas aeruginosa. Genetics, 186(4):1345–1354.

Manioudaki, M. E. and Poirazi, P. (2013). Modeling regulatory cascades using Artificial Neural Networks: the case of transcriptional regulatory networks shaped during the yeast stress response. Frontiers in genetics, 4:110.

McEntire, K. D., Gage, M., Gawne, R., Hadfield, M. G., Hulshof, C., Johnson, M. A., Levesque, D. L., Segura, J., and Pinter-Wollman, N. (2021). Understanding drivers of variation and predicting variability across levels of biological organization. Integrative and Comparative Biology, 61(6):2119–2131. Publisher: Oxford University Press.

Mousseau, T. A. and Roff, D. A. (1987). Natural selection and the heritability of fitness components. Heredity, 59(2):181–197.

Nijhout, H. F., Berg, A. M., and Gibson, W. T. (2003). A mechanistic study of evolvability using the mitogen-activated protein kinase cascade. Evolution & Development, 5(3):281–294.

Nijhout, H. F. and Reed, M. C. (2014). Homeostasis and dynamic stability of the phenotype link robustness and plasticity. American Zoologist, 54(2):264–275.

Orgogozo, V., Morizot, B., and Martin, A. (2015). The differential view of genotype–phenotype relationships. Frontiers in genetics, 6:179.

Paschold, A., Marcon, C., Hoecker, N., and Hochholdinger, F. (2010). Molecular dissection of heterosis manifestation during early maize root development. Theor Appl Genet, 120(2):383–388.

Perry, M. W., Bothma, J. P., Luu, R. D., and Levine, M. (2012). Precision of hunchback expression in the Drosophila embryo. Current biology, 22(23):2247–2252.

Petrizzelli, M., Coton, C., and de Vienne, D. (2024). Formalizing the law of diminishing returns in metabolic networks using an electrical analogy. R. Soc. Open Sci., 11(10):240165.

Powell, J. E., Henders, A. K., McRae, A. F., Kim, J., Hemani, G., Martin, N. G., Dermitzakis, E. T., Gibson, G., Montgomery, G. W., and Visscher, P. M. (2013). Congruence of additive and non-additive effects on gene expression estimated from pedigree and SNP data. PLoS genetics, 9(5):e1003502.

Pryciak, P. M. (2008). Customized Signaling Circuits. Science, 319(5869):1489–1490.

Radde, N. E. and Hütt, M.-T. (2016). The Physics behind Systems Biology. EPJ Nonlinear Biomed Phys, 4(1):7.

Rest, J. S., Morales, C. M., Waldron, J. B., Opulente, D. A., Fisher, J., Moon, S., Bullaughey, K., Carey, L. B., and Dedousis, D. (2013). Nonlinear Fitness Consequences of Variation in Expression Level of a Eukaryotic Gene. Molecular Biology and Evolution, 30(2):448–456.

Roff, D. A. and Mousseau, T. A. (1987). Quantitative genetics and fitness: lessons from Drosophila. Heredity, 58(1):103–118.

Rosas, U., Barton, N. H., Copsey, L., Barbier de Reuille, P., and Coen, E. (2010). Cryptic variation between species and the basis of hybrid performance. PLoS biology, 8(7):e1000429.

Rossignol, R., Faustin, B., Rocher, C., Malgat, M., Mazat, J. P., and Letellier, T. (2003). Mitochondrial threshold effects. Biochemical Journal, 370:751–762.

Simon, H. A. (1962). The architecture of complexity. Proceedings of the American Philosophical Society, 106:467–482.

Snoep, J. L., Yomano, L. P., Westerhoff, H. V., and Ingram, L. O. (1995). Protein burden in Zymomonas mobilis: negative flux and growth control due to overproduction of glycolytic enzymes. Microbiology, 141(9):2329–2337.

Szathmáry, E. and Maynard Smith, J. (1995). The major evolutionary transitions. Nature, 374(6519):227–232. Number: 6519 Publisher: Nature Publishing Group.

Tokuriki, N., Jackson, C. J., Afriat-Jurnou, L., Wyganowski, K. T., Tang, R., and Tawfik, D. S. (2012). Diminishing returns and tradeoffs constrain the laboratory optimization of an enzyme. Nature Communications, 3(1):1257.

Tokuriki, N. and Tawfik, D. S. (2009). Protein Dynamism and Evolvability. Science, 324(5924):203–207.

Umerez, J. (2016). Biological Organization from a Hierarchical Perspective. Evolutionary theory: A hierarchical perspective. University of Chicago Press.

Veitia, R. A. (2003). A sigmoidal transcriptional response: cooperativity, synergy and dosage effects. Biological Reviews, 78(1):149–170.

Vilaprinyo, E., Alves, R., and Sorribas, A. (2010). Minimization of biosynthetic costs in adaptive gene expression responses of yeast to environmental changes. PLoS Comput Biol, 6(2):e1000674.

Violle, C., Navas, M., Vile, D., Kazakou, E., Fortunel, C., Hummel, I., and Garnier, E. (2007). Let the concept of trait be functional! Oikos, 116(5):882–892.

Visscher, P. M., Hill, W. G., and Wray, N. R. (2008). Heritability in the genomics era—concepts and misconceptions. Nature reviews genetics, 9(4):255–266.

Whitacre, J. M. (2012). Biological robustness: paradigms, mechanisms, and systems principles. Front. Genet., 3:67.

Wright, S. (1934). Physiological and evolutionary theories of dominance. American Naturalist, 68:24–53.

Xing, J., Sun, Q., and Ni, Z. (2016). Proteomic patterns associated with heterosis. Biochimica et Biophysica Acta (BBA)-Proteins and Proteomics, 1864(8):908–915.

Yang, R.-C. (2017). Genome-wide estimation of heritability and its functional components for flowering, defense, ionomics, and developmental traits in a geographically diverse population of Arabidopsis thaliana. Genome, 60(7):572–580.

Zhang, J. (2018). Neutral theory and phenotypic evolution. Molecular biology and evolution, 35(6):1327–1331.

Zhou, P., Hirsch, C. N., Briggs, S. P., and Springer, N. M. (2019). Dynamic patterns of gene expression additivity and regulatory variation throughout maize development. Molecular plant, 12(3):410–425.

